# Time-course analysis of early meiotic prophase events informs mechanisms of homolog pairing and synapsis in *Caenorhabditis elegans*

**DOI:** 10.1101/143693

**Authors:** Susanna Mlynarczyk-Evans, Anne M Villeneuve

**Author notes:** Corresponding Author: Anne M. Villeneuve 279 Campus Dr. B300 Beckman Center Stanford University School of Medicine Stanford, CA 94305.

## Abstract

Segregation of homologous chromosomes during meiosis depends on their ability to reorganize within the nucleus, discriminate among potential partners, and stabilize pairwise associations through assembly of the synaptonemal complex (SC). Here we report a high-resolution time-course analysis of these key early events during *Caenorhabditis elegans* meiosis. Labeled nucleotides are incorporated specifically into the X chromosomes during the last two hours of S phase, a property we exploit to identify a highly synchronous cohort of nuclei. By tracking X-labeled nuclei through early meiotic prophase, we define the sequence and duration of chromosome movement, nuclear reorganization, pairing at pairing centers (PCs), and SC assembly. Appearance of ZYG-12 foci (marking attachment of PCs to the nuclear envelope) and onset of active mobilization occur within an hour after S phase completion. Movement occurs for nearly 2 hours before stable pairing is observed at PCs, and autosome movement continues for roughly 4 hours thereafter. Chromosomes are tightly clustered during a 2-3 hour post-pairing window, during which the bulk of SC assembly occurs; however, initiation of SC assembly can precede evident chromosome clustering. SC assembly on autosomes begins immediately after PC pairing is detected and is completed within about 3.5 hours. For the X chromosomes, PC pairing is contemporaneous with autosomal pairing, but autosomes complete synapsis earlier (on average) than X chromosomes, implying that X chromosomes have a delay in onset and/or a slower rate of SC assembly. Additional evidence suggests that transient association among chromosomes sharing the same PC protein may contribute to partner discrimination.

## Introduction

It is well known that the germ line of the nematode *Caenorhabditis elegans* is organized in a spatio-temporal gradient of germ cells entering and progressing through meiotic prophase along the distal-proximal axis of the gonad (Hillers *et al*. 2015). This property has been a major factor in the emergence of *C. elegans* as an important model system for investigating meiotic mechanisms, as it enables pseudo-time-course analyses of meiotic events, which are conducted by quantifying specific nuclear and chromosomal features in spatially-defined zones or rows of nuclei distributed along the gonad axis (Colaiacovo *et al*. 2003; Macqueen *et al*. 2002). While this has been and continues to be a powerful approach, there are inherent limitations because nuclei that enter meiosis contemporaneously are distributed over several rows within the gonad ((Mlynarczyk-Evans *et al*. 2013) and this work), resulting in local staging heterogeneity.

Here, we use the complementary approach of S phase labeling of chromosomal DNA to follow progression of meiotic prophase germ cells. Previously, a more coarse-grained time course analysis used S phase labeling to estimate the timing of progression through the entirety of meiotic prophase (Jaramillo-Lambert *et al*. 2007). That study revealed that duration of meiotic prophase during oogenesis in *C. elegans* hermaphrodites is on the order of 2-3 days, and varies depending on age and presence/absence of sperm. In the work presented here, we have conducted a more fine-grained time course analysis focusing on the early events in meiotic prophase, exploiting the late-replicating property of the X chromosomes(Jaramillo-Lambert *et al*. 2007; Mlynarczyk-Evans *et al*. 2013) to identify a highly synchronous cohort of nuclei that were in late S phase at the time that label was introduced. By analyzing multiple features of the meiotic program at time points distributed through the first 8 hours post labeling, we have elucidated the timing of onset and duration of key early meiotic prophase events and the temporal relationships among them. We further demonstrate that quantifying chromosomal features in temporally-defined nuclear cohorts has the capacity to reveal aspects of the meiotic program not previously detected by other approaches.

## Materials and Methods

All experiments (except those shown in Figure 3) were conducted using wild type *C. elegans* Bristol N2 hermaphrodite worms cultivated at 20°C under standard conditions.

### S-phase Labeling

S-phase labeling was performed by microinjection of fluorescent nucleotides as described in (Hayashi *et al*. 2010), except for Figure 5 experiments involving ZIM-2, where S-phase labeling was accomplished by feeding with EdU-labeled bacteria for 5 hours as described in (Rosu *et al*. 2011). Comparable X-specific labeling and progression of nuclei with labeled X chromosomes through meiosis, including similar spatial distribution of labeled nuclei, were observed with both labeling methods. For injection experiments, gonads of young adult worms (24 hr post L4) were microinjected with 0.1 mM Cy3-dCTP (GE Health Sciences), and worms were recovered and maintained on food at 20°C until they were dissected, fixed and processed for immunoflourescence at the indicated time points post-injection. Gonads were injected in a position midway between the distal tip and the turn, thereby avoiding and minimizing potential mechanical disruption of the region of the gonad in which meiotic features would later be assessed. For two-color S-phase labeling experiments, injection of Cy3-dCTP was followed by a second injection of FITC-labeled dNTPs at the indicated times (Figure 1E). For Figure 5 ZIM-2 experiments, EdU incorporation was detected using click chemistry as in (Rosu *et al*. 2011).

**Figure 1.**
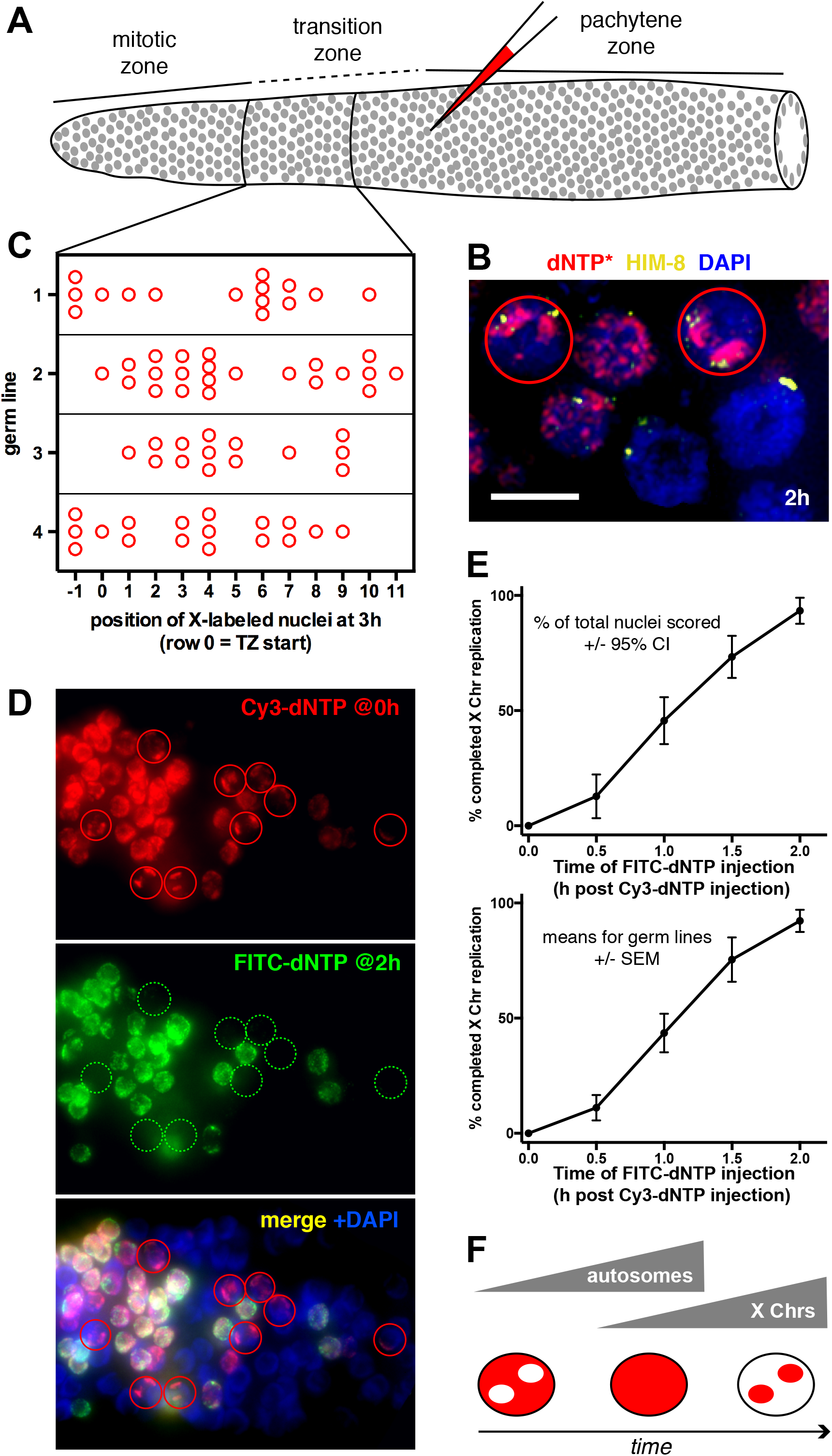
Timing of X Chromosome replication during meiotic S phase. (A) Schematic diagram of a hermaphrodite gonad being injected with fluorescently-labelled dNTPs (red); gray circles indicate germ cell nuclei. Black lines flank the region where nuclei that had incorporated fluorescent nucleotides specifically into their X chromosomes were located at 3 h post injection. (B) Field of nuclei from an injected gonad, with nuclei in which labelled nucleotides were incorporated specifically into the X chromosomes (indicating that they were in late S phase at the time of injection) circled in red; immunostaining of the X-chromosome pairing center binding protein HIM-8 is used to identify the X chromosomes. The field also includes nuclei that were not in S phase (no red label) or that were in mid S phase (red label distributed through the nucleus) at the time of injection. Image is a projection of a 3D data stack encompassing whole nuclei; scale bar = 5 μm. (C) Diagram showing the spatial distribution (along the distal-proximal axis) of nuclei with X-specific labelling within 4 example germ lines; such nuclei were typically distributed over 9-12 rows. “TZ start” was defined as the most distal row containing two or more nuclei displaying highly clustered chromosomes. (D) Example images from a series doublelabelling experiment designed to measure the duration of the period of exclusive X chromosome replication (orientation: L to R = distal to proximal). In the images shown, FITC-labelled dNTPs (green) were injected 2 h after injection of Cy3-labelled dNTPs (red). Nuclei with X-specific incorporation of Cy3-dNTP are circled; although FITC-dNTP was readily incorporated into nuclei with Cy3-labelled autosomes, it was not incorporated into nuclei with X-specific Cy3 labelling, indicating that such nuclei had completed replication in the interval between injections. (E) Graphs of time course analysis of X chromosome replication. The x axis indicates the time interval between Cy3-dNTP injection and FITC-dNTP injection. The y axis indicates the % of nuclei in the X-specific Cy3-labelled cohort that did NOT incorporate FITC-dNTP when it was injected at the indicated time post Cy3-dNTP injection, reflecting completion of X replication prior to the second injection. In the top graph, data are presented as “percent of total nuclei with X-specific Cy3 labelling” (± 95% CI), whereas the bottom graph plots the means of the percentages scored for individual germ lines (± SEM). Number of gonads/ nuclei scored for each time point (h post Cy3-dNTP injection) were as follows: 0 h, 5 gonads/79 nuclei; 0.5 h, 5 gonads/47 nuclei; 1.0 h, 6 gonads/92 nuclei; 1.5 h, 5 gonads/90 nuclei; 2.0 h, 5 gonads/75 nuclei. (F) Schematic illustrating progression of autosomal and X-chromosome replication during S-phase and three distinct cytological phenotypes that can be observed for S-phase single-labeling experiments: Early S phase (labeled autosomes, unlabeled X chromosomes), Mid S phase (autosomes and X chromosomes labeled), and Late S phase (only X chromosome labeled).

### Immunofluorescence

Dissection of gonads, fixation, immunostaining, and DAPI staining were performed essentially as in (Martinez-Perez and Villeneuve 2005). The following primary antibodies were used: guinea pig anti-HIM-8 (1:500 (Phillips *et al*. 2005)); rabbit anti-ZIM-3 (1:2,000 (Phillips and Dernburg 2006)); chicken anti-HTP-3 (1:250 ((Macqueen *et al*. 2005)); rabbit anti-SYP-1 (1:250 (Macqueen *et al*. 2002)). Secondary antibodies were Alexa Fluor 488, 555, and/or 647-conjugated goat antibodies directed against the appropriate species (1:400; Invitrogen).

### Image Collection, Rendering and Analysis

For fixed specimens, 3-D images were collected as Z-stacks (0.1 or 0.2 μm step size) using a 60X NA 1.42 objective with 1.5X optivar on a DeltaVision widefield deconvolution microscopy system (Applied Precision) and deconvolved using softWoRx software. Images in Figure 1 are maximum intensity projections generated in softWoRx. For images of individual nuclei in Figures 2 and 4-7, contrast adjustments, 3-D cropping and image rendering (2-D maximum intensity projections or 3-D surface opacity renderings) were performed using the Volocity 5 software package (PerkinElmer). Inter-focus distances reported in Figure 5 were measured using Volocity software.

**Figure 2.**
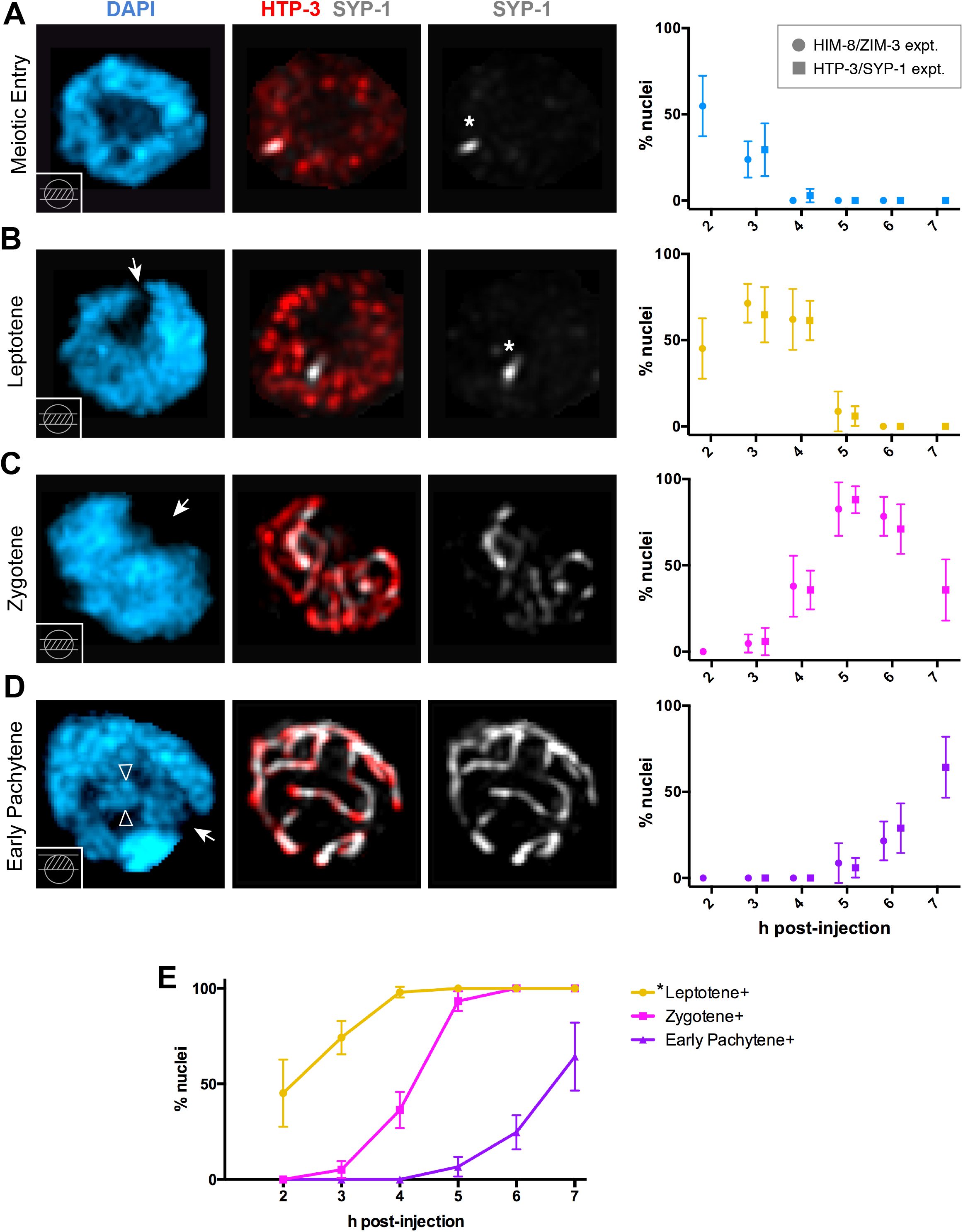
Distinct organization of DAPI-stained chromatin during early meiotic prophase substages allows alignment of data from different experiments. (A-D left) Immunofluorescence images of individual nuclei stained for DNA(DAPI), meiotic chromosome axis protein HTP-3, and SC central region protein SYP-1, illustrating the DAPI organization features used to identify and define the indicated substages; images are maximum intensity projections generated in Volocity, with the portion of the Z-stack projected depicted in the inset diagrams at the lower left of the DAPI panels. Arrows pointing to regions lacking fluorescence signal in the DAPI panels for B-D indicate the inferred position of the nucleolus; asterisks in A and B indicate a single aggregate of SYP proteins present in meiotic prophase nuclei prior to initiation of SC assembly; paired arrowheads in (D) indicate parallel tracks of DAPI staining reflecting aligned and synapsed homologs. Whereas the nucleus depicted in B is clearly at the leptotene stage, we have used the “Leptotene*” designation for this DAPI category as some nuclei with a similar DAPI configuration exhibit a very short stretch of axis-associated SYP-1 in addition to a SYP-1 aggregate indicating that this DAPI category also includes nuclei transitioning to early zygonema; see main text and Figure 6. (A-D right) Graphs depicting the percentages of X-specific S-phase labeled nuclei exhibiting the DAPI configuration characteristic of the illustrated stage at the indicated time after injection (± 95% CI). Data from two different immunofluorescence experiments show strong concordance. Number of gonads/ nuclei scored for each time point (h post Cy3-dNTP injection) for the HIM-8/ZIM-3 time course were as follows: 2h, 3 gonads/31 nuclei; 3h, 5 gonads/63 nuclei; 4h, 5 gonads/58 nuclei; 5h, 4 gonads/23 nuclei; 6h, 4gonads/51 nuclei. Number of gonads/ nuclei scored for each time point for the HTP-3/SYP-1 time course were as follows: 3h, 3 gonads/34 nuclei; 4h, 6 gonads/70 nuclei; 5h, 6gonads/67 nuclei; 6h, 6 gonads 38 nuclei; 7h 4 gonads/28 nuclei. (E) Cumulative distribution “Event curves” plotting the percent of nuclei that had reached or passed the indicated stage (defined by DAPI morphology); data from the two different types of experiments plotted in the A-D graphs were combined. The “Leptotene*” designation indicates that this category also includes nuclei transitioning to early zygonema.

### Live “*ex vivo*” Imaging

For live imaging of ZYG-12::GFP in S-phase labeled nuclei, Cy3-dCTP was injected into hermaphrodites of strain WH223 (*ojIs9 [zyg-12ABC::gfp unc-119(+)]; unc-119(ed3) III*) (Malone *et al*. 2003). Imaging was carried out on partially extruded gonads, as in (Zhang *et al*. 2012), with modifications. Worms were dissected in 10 μl of M9 containing tricaine (0.1%) and tetramisole (0.01%), and the coverslip with the dissected worms was flipped onto a 2% agarose pad and imaged immediately using the DeltaVision deconvolution microscopy system with a 60x oil objective with 1.5x optivar. Movies were acquired as stacks of 3 optical sections (1.5 μm step size) every 10 s over a 2 min time period. After acquisition of movies to verify movement of ZYG-12::GFP patches, 3D image stacks (0.2 μm step size) encompassing the full depth of nuclei were also acquired and deconvolved to enable classification of nuclei based on ZYG-12::GFP aggregate patterns.

Statistical analyses (two-tailed Fisher exact tests) were conducted using GraphPad Prism or Graphpad QuickCalcs software.

All relevant data (and representative images) are presented within the paper; complete imaging data sets are available upon request. Nematode strains used are available from the Caenorhabditis Genetics Center (funded by NIH Office of Research Infrastructure Programs P40 OD010440).

## Results and Discussion

### X-specific S-phase labeling is distributed over 9-12 rows of germ cell nuclei

To mark nuclei by incorporation of fluorescently labeled nucleotides during the S phase preceding meiotic prophase, Cy3-labeled dNTPs were injected into a common central cytoplasmic core (the rachis) that is connected to germ cells *via* cytoplasmic bridges (Figure 1A). When gonads were subsequently analyzed using fluorescence microscopy, a subset of labeled nuclei had incorporated Cy3-labeled nucleotides specifically into their X chromosomes, which replicate out of phase with the autosomes, as previously reported ((Jaramillo-Lambert *et al*. 2007), Figure 1B). We found that these nuclei with X-specific Cy3 labeling, which comprise a highly synchronous cohort that were in late S phase at the time of label incorporation, were typically distributed over a spatial domain of 9 -12 rows within germ lines that were analyzed at 2-3 h after injection (Figure 1C and (Mlynarczyk-Evans *et al*. 2013)). This relatively broad spatial distribution of temporally synchronous nuclei represents a potential confounding factor for pseudo-time course analyses in which meiotic features are quantified in spatially-defined zones along the gonad axis. Thus, time course analyses based on S-phase labeling should have improved power to detect transient nuclear or chromosomal features and/or temporal relationships among different features.

### The period of exclusive X replication lasts approximately two hours

To determine the length of the period when only the X chromosomes are replicating, we conducted two-color labeling experiments in which we followed injection of Cy3-labeled dNTPs with a second injection of FITC-labeled dNTPs after several different time intervals and determined the fraction of nuclei with X-specific Cy3 incorporation that had also incorporated FITC-labeled nucleotides (Figure 1D,E). Incorporation of the second label indicated that S phase was still ongoing at the time when the second label was introduced, whereas failure to incorporate the second label indicated that replication of the X chromosomes was completed during the interval between injection of the first and second label. We found that when the two injections were separated by a 2 hour interval, essentially all nuclei showing X-specific incorporation of Cy3 lacked the FITC label (Figure 1D,E), indicating that these nuclei had completed replication within that 2 hour interval. These data indicate that the period of exclusive X replication lasts between 1.5 and 2 hours.

In our injection experiments, we noted two additional cytologically distinct classes of nuclei: 1) those in which Cy3 label was detected in all chromosomes, corresponding to nuclei that were in mid-S phase at the time of injection, and 2) those in which Cy3 label was detected in the autosomes but lacking in the X chromosomes, corresponding to nuclei that were in early S phase at the time of injection. Except where noted, in all of the experiments presented in the remainder of this manuscript, the indicated features were assessed specifically in the cohort of nuclei exhibiting X-specific incorporation of labeled nucleotides (following a single injection), *i.e*., the scored nuclei had been in the last two hours of S phase at the time of injection.

### Distinct organization of DAPI-stained chromatin during early meiotic prophase substages allows alignment of data from different experiments

A major goal of this work was to reveal the temporal relationships among multiple distinct events occurring during early meiotic prophase. Since it is not possible to assess all relevant features in the same experiment, it is essential to be able to combine and overlay the data from different experiments. To this end, we evaluated the organization of DAPI-stained chromatin in nuclei from two separate time course experiments that were designed to evaluate different features of meiosis (Figure 2). Specifically, we examined images of DAPI-stained chromatin from: 1) a time course in which we assessed pairing at the pairing centers (PCs) of the X chromosomes (marked by HIM-8 (Phillips *et al*. 2005)) and at the PCs of chromosomes I and IV (marked by ZIM-3 (Phillips and Dernburg 2006)); and 2) a time course experiment in which we assessed the status of meiotic chromosome axes (visualized by immunostaining of HORMA domain protein HTP-3 (Goodyer *et al*. 2008; Macqueen *et al*. 2005)) and assembly of the synaptonemal complex (SC, visualized by immunostaining for SC central region protein SYP-1 (Macqueen *et al*. 2002)). Importantly, we sorted nuclei into four classes based on examination of DAPI images alone (Figure 2A-D left; see below)), and for both time courses, we plotted the frequencies of X-labeled nuclei in each DAPI class as a function of time (Figure 2A-D right). The data from the two different types of experiments showed strong concordance, indicating that alignment of separate time course experiments is justified and can be used to infer the relative timing of features measured in different experiments.

While we will defer a more extensive and detailed analysis of the timing of SC assembly to a later section, we will discuss here how the combination of X-specific S-phase labeling, DAPI staining, and immunostaining of HTP-3 and SYP-1 enabled us to refine our identification of different sub-stages of meiotic prophase:

First, we could differentiate distinct “meiotic entry” and leptotene stages based on the relative position of the chromatin and the nucleolus. In both types of nuclei, the overall shape of the territory occupied by the DAPI-stained chromatin was roughly spherical, parallel pairs of DAPI tracks were not observed, HTP-3 was in the process of coalescing into axial structures, and in most cases (but see below), SYP-1 protein was visible but limited to a single nuclear aggregate. However, such nuclei fell into two classes: 1) nuclei with the DAPI-dark nucleolus located in the center of the nucleus, and 2) nuclei in which the DAPI-dark nucleolus had adopted an overtly off-center position (sometimes with a small DAPI-dark region adjacent to the nuclear envelope [NE]). We use the term “Meiotic Entry” to refer to the former class (which likely includes nuclei in the early leptotene stage), as such nuclei declined in abundance concomitant with an increase in the abundance of the latter class, which is considered to correspond predominantly to the (mid-to-late) leptotene stage.

The leptotene stage transitioned into the zygotene stage, during which SC assembly occurs. The bulk of zygotene nuclei are characterized by well-defined HTP-3 axes with SYP-1 localized to a subset of these, some parallel DAPI tracks, a markedly non-spherical DAPI-stained chromatin mass and a highly off-center nucleolus. Zygonema gave way to early pachynema, defined by localization of SYP-1 along the full lengths of the HTP-3 axes, extensive parallel tracks of DAPI-stained chromatin, and return to a more spherical DAPI-stained chromatin mass and more centrally-positioned nucleolus, albeit with a smaller chromosome-free region still remaining adjacent to the NE.

Data from several different experiments were combined to generate the “Event curves” shown in Figure 2E. This graph plots cumulative distribution curves indicating the fraction of (X-specific labeled) nuclei that had reached or completed that stage at the indicated time points post injection. These event curves allow us to make inferences regarding the duration of the indicated stages as defined by DAPI morphology. Given the approximately 2 h window of exclusive incorporation of labeled nucleotides into the X chromosomes, we infer that approximately 50% of X-labeled nuclei will have completed S phase by 1 h post injection. Thus, the 2 h post injection time point corresponds to 1 h after completion of S phase. Similarly, as 50% of nuclei have acquired the “leptotene” DAPI morphology by that time, we infer that the leptotene stage begins within about 1 h after completion of S phase; however, as a few short stretches of SC are detected in a subset of nuclei with this morphology (see below), we use the “*Leptotene” designation in the graph to indicate that this DAPI category also includes nuclei in the early zygotene stage. Acquisition of the characteristic zygotene clustered chromatin morphology begins about 3 hours after S phase completion and lasts about 2.5 - 3 hours. Finally, the early pachytene stage begins between 5.5 - 6 hours after S phase completion. These event curves will serve as a frame of reference for the different analyses discussed below.

### Timing of formation and dissolution of Pairing Center-associated mobile ZYG-12 aggregates

The reorganization of chromosomes within meiotic prophase nuclei in *C. elegans* is driven by the attachment of specialized chromosome regions known as pairing centers (PCs) to the cytoskeletal motility apparatus through association with a conserved NE-bridging complex composed of a SUN-domain protein (SUN-1) and a KASH-domain protein (ZYG-12) (Woglar and Jantsch 2014). Although the nature of these PC-mediated movements on time scales of seconds and minutes has been evaluated using germ cells expressing GFP::SUN-1 or ZYG-12::GFP (Baudrimont *et al*. 2010; Wynne *et al*. 2012), the duration of the movement-competent state has not been reported.

To evaluate the duration of the stage(s) during which PC-mediated chromosome movement occurs, we used live imaging to assess X-labeled cohorts of nuclei for the appearance, pattern/organization, and disappearance of mobile nuclear-envelope associated aggregates of ZYG-12::GFP (Figure 3; Movies S1-2). These ZYG-12::GFP foci or patches represent functional attachments between PCs on the chromosomes, through the NE-spanning SUN-KASH complexes, to the dynein motors that mediate chromosome movement (Penkner *et al*. 2007; Sato *et al*. 2009).

**Figure 3.**
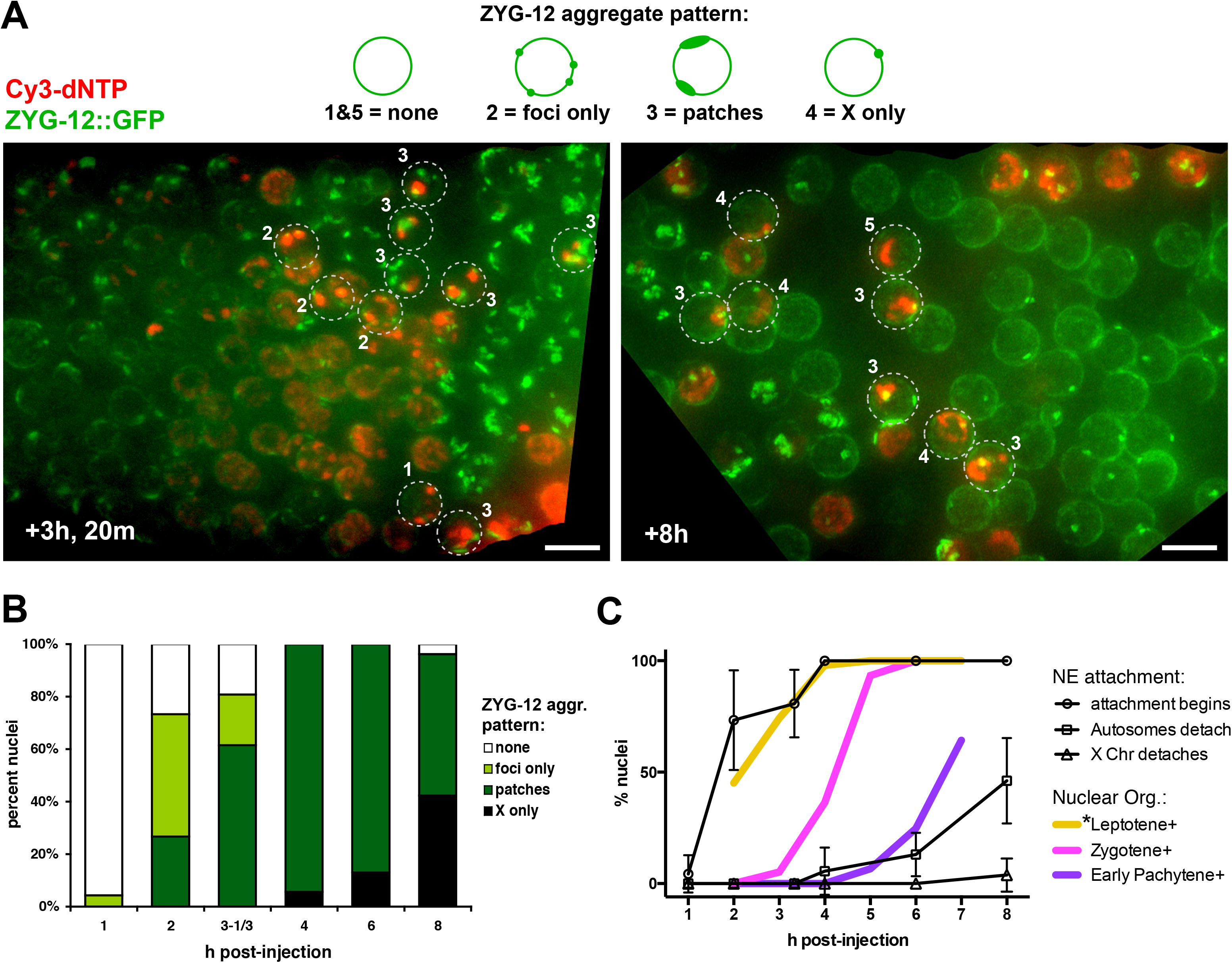
Timing of ZYG-12 aggregate formation and disappearance. (A) Live images of Cy3-labelled chromosomes and ZYG-12::GFP in fields of germ cell nuclei from partially extruded gonads imaged at the indicated times post injection (orientation: L to R = distal to proximal). Images are projections of deconvolved 3D stacks taken after collection of live movies which verified that whenever ZYG-12::GFP foci or patches were observed, they were moving (see Materials and Methods). Nuclei with X-specific Cy3 label are indicated by dashed circles; nuclei were classified based on the ZYG-12 aggregate patterns indicated in the schematics above the images. Scale bar = 5 μm. (B) Stacked bar graph indicating the distribution of X-labeled nuclei among the ZYG-12 aggregate pattern classes at the indicated times post injection. Number of gonads/ nuclei scored for each time point (h post Cy3-dNTP injection) were: 1h, 4 gonads/23 nuclei; 2h, 4 gonads/15 nuclei; 3.33h, 3 gonads/25 nuclei; 4h, 3 gonads/18 nuclei; 6h, 5 gonads/46 nuclei; 8 h, 4 gonads/26 nuclei. (C) Cumulative distribution “Event curves”, using ZYG-12 aggregate patterns as indicators of functional attachment of autosomal and X chromosome pairing centers to the NE (± 95% CI) (see text).

Our analysis indicated that mobile NE attachments can be detected within an hour after completion of S phase; ZYG-12 foci or patches are observed to be moving as soon as they are detected. ZYG-12 aggregates corresponding to autosomal pairing centers remain visible and active for about 6 h thereafter. Thus, PC movement precedes acquisition of the clustered chromosome organization characteristic of the zygotene stage by more than two hours, and autosomal PCs remain active for about 1.5 hours after nuclei reach the early pachytene stage, and then become inactive. As previously reported, the X-chromosome PCs remain active beyond the time when the autosomal PCs have lost their ZYG-12 aggregates (Penkner *et al*. 2007; Sato *et al*. 2009); however, our experiments did not address the duration of the stage of active X-PC movement, as the X-associated ZYG-12 aggregates persisted past the time window analyzed in our experiments.

### Pairing at X and autosome PCs occurs contemporaneously within a narrow time window

We used immunostaining for HIM-8, which localizes at the X-PCs, and ZIM-3, which localizes to the PCs of chromosomes I and IV, to assess the timing of pairing at the PCs (Figure 4). Plots of the frequencies of pairing at the X-PC and at the PCs of chromosomes I and IV at different times post injection were largely superimposable (Figure 4B). Further, at all time points analyzed, the three assayed PCs were either all paired or all unpaired in the vast majority of nuclei (Figure 4C). In nuclei in which only a subset of the assayed PCs were paired, the observed classes of nuclei (Figure 4A, C) were consistent with previous reports indicating no apparent temporal hierarchy for pairing of X chromosomes vs. autosomes (Macqueen *et al*. 2002; Nabeshima *et al*. 2011)). Placed in the context with our data for other features analyzed above (Figure 4D), our data indicate that pairing at the 3 PCs assayed in our experiments occurs predominantly within a narrow time window of roughly 30 minutes, beginning approximately 2 hours after the onset of ZYG-12-mediated PC attachment and movement, and that the X chromosomes and autosomes pair contemporaneously within the same time window; the true pairing window for the full cohort of six chromosome pairs is likely to be somewhat broader. Further, completion of pairing at PCs coincides roughly with acquisition of the characteristic zygotene nuclear organization.

**Figure 4.**
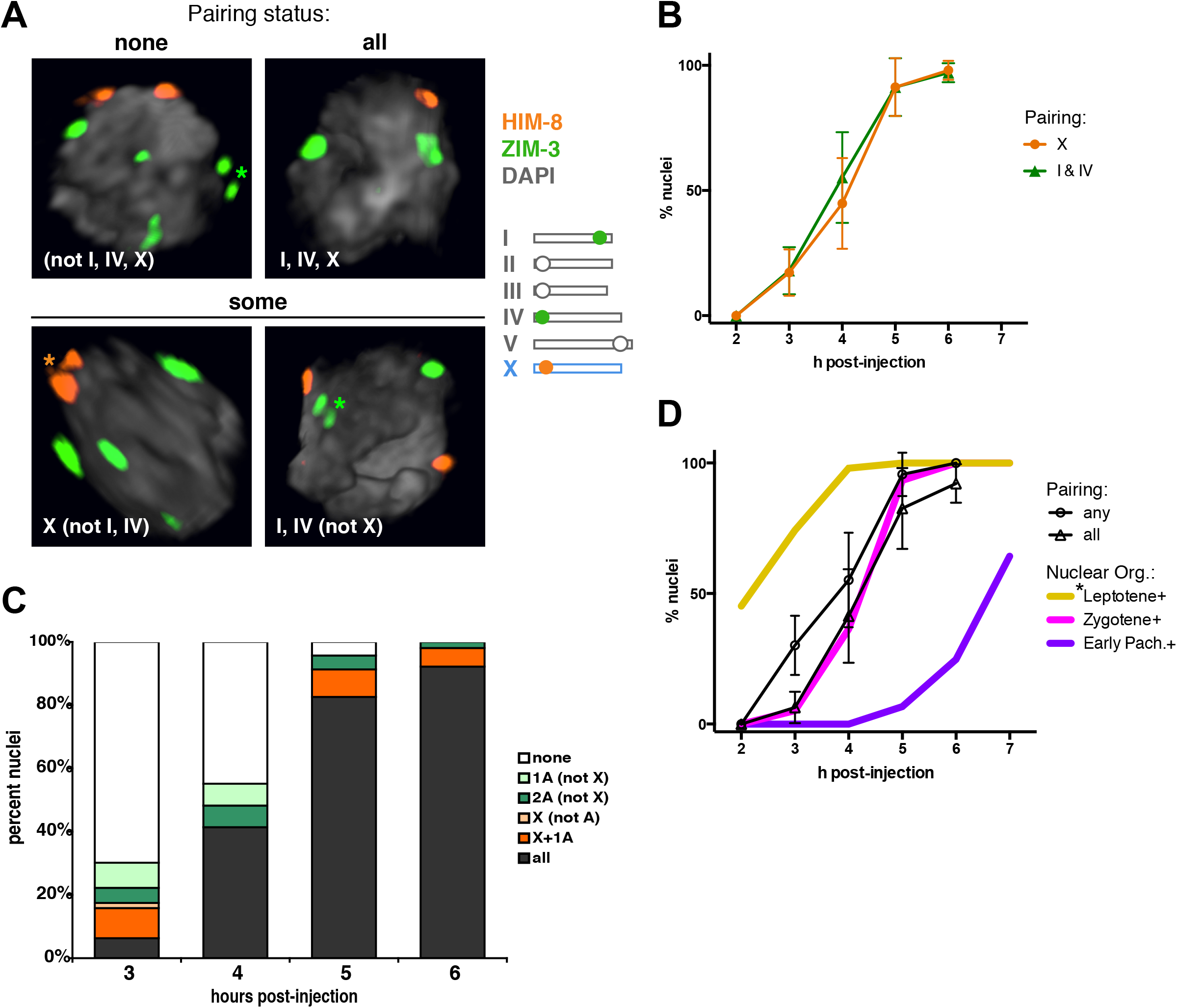
Timing of pairing at X and autosome pairing centers. (A) Images of early prophase nuclei immunostained for HIM-8 (the X-PC binding protein) and ZIM-3 (which associates with the PCs of chromosomes I and IV), showing examples where none of the three PCs are paired, all three are paired, and some of the three PCs are paired (only X, or only I and IV). Images shown are surface renderings of DAPI-stained chromatin and maximum intensity projections of immunofluorescence signals. Both before and after pairing, the PC protein signals sometimes appear as a group of 2-3 close but resolvable signals (asterisks) perhaps due to the fact that there are several distinct clusters of ZIM-3 and HIM-8 binding motifs on chromosomes I and X, respectively (Nabeshima *et al*. 2011; Phillips *et al*. 2009), or due to stretching forces exerted at the PC during chromosome movement. (B) Graph plotting the percent of X-labeled nuclei exhibiting pairing at the X-PCs (orange) or at the PCs of both I and IV (green); error bars indicate 95% CI. Number of gonads/nuclei scored for each time point were: 2h, 3 gonads/32 nuclei; 3h, 5 gonads/63 nuclei; 4h, 5 gonads/29 nuclei; 5h, 4 gonads/23 nuclei; 6h, 4 gonads/51 nuclei. (C) Stacked bar graphs showing frequencies observed for each of the indicated pairing configurations. (D) Graph plotting the percent of X-labeled nuclei exhibiting pairing at the PCs of “any” or “all” of the assayed chromosomes, together with the cumulative distribution event curves for nuclear organization. Together, the data indicate that pairing at the assayed PCs occurs within a narrow time window of roughly 30 minutes, beginning approximately 2 hours after the onset of ZYG-12-mediated PC attachment and movement, and that the X chromosomes and autosomes pair within the same time window.

### PCs of chromosomes that share the same PC binding protein exhibit preferential proximity during the zygotene stage

Whereas the X-PC specifically recruits HIM-8 and the V-PC specifically recruits ZIM-2, chromosomes I and IV share the same PC-binding protein (ZIM-3), as do chromosomes II and III (ZIM-1) (Phillips and Dernburg 2006; Phillips *et al*. 2005). This finding had naturally raised the question of whether the PCs of chromosomes that share the same PC binding protein might tend to associate with each other more frequently that those that do not. Evidence for such preferential association was not detected in a previous study, but staging heterogeneity in spatially-defined gonad zones may have limited the sensitivity of that prior analysis. Our time course analysis provided an opportunity to revisit this issue by measuring distances between the PCs of different chromosomes in a temporally defined cohort of meiotic prophase nuclei (Figure 5).

**Figure 5.**
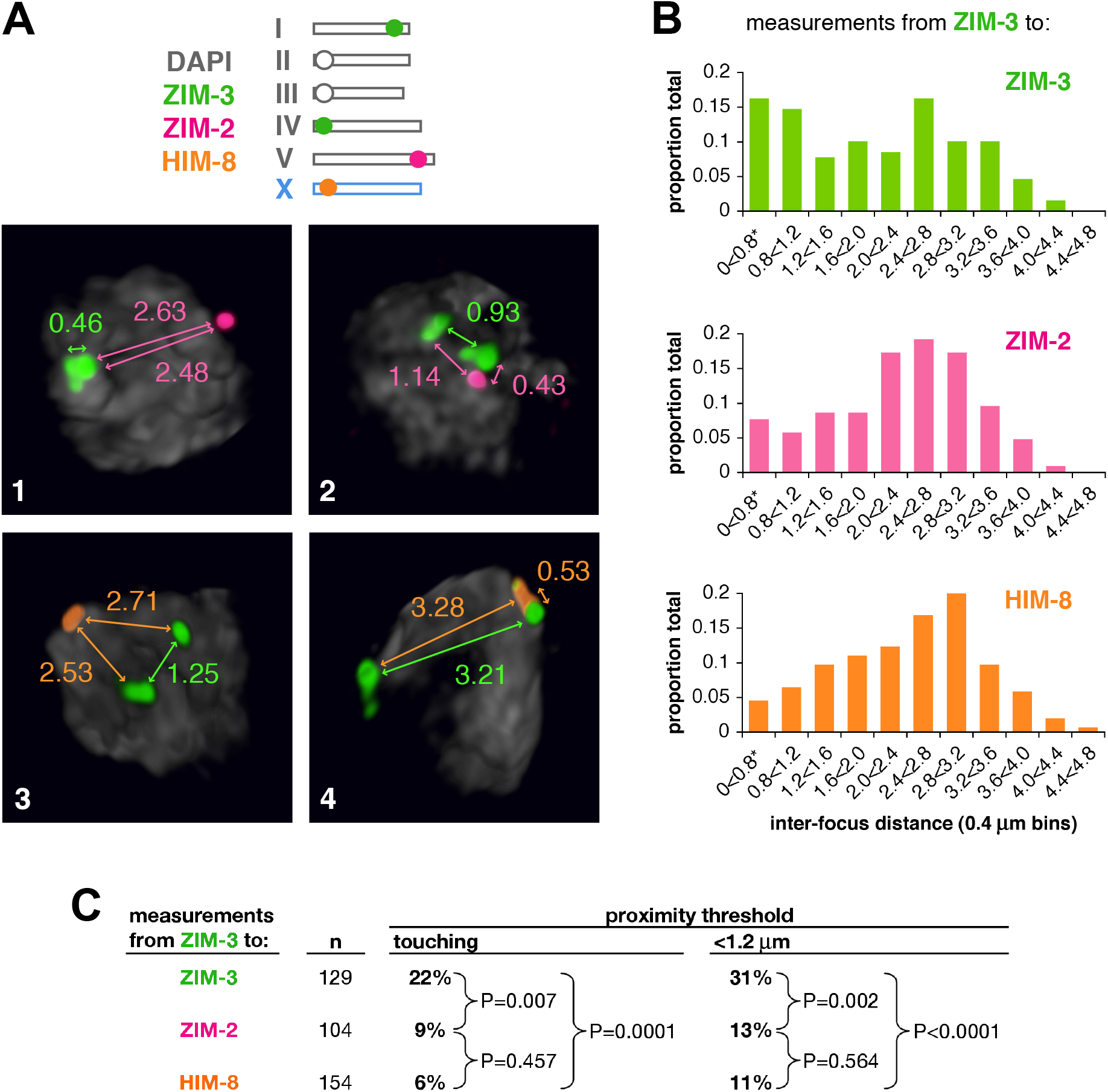
PCs of chromosomes that share the same PC binding protein exhibit preferential proximity during the zygotene stage. (A) Images of nuclei costained for either ZIM-3 (PC-binding protein for chromosomes I and IV, as indicated in the schematic) and HIM-8 (X-PC binding protein) or ZIM-3 and ZIM-2 (PC-binding protein for chromosome V). In all nuclei scored in this analysis (taken from the 4, 5 and 6 h post injection time points, *i.e*., 3-5 h post S phase), the PCs of the homologs were already paired, as indicated by a single HIM-8 or ZIM-2 focus and 2 separate (or immediately adjacent) ZIM-3 foci. For the ZIM-3 and HIM-8 analysis, number of gonads/nuclei scored for each time point were: 4 h, 5 gonads/12 nuclei; 5 h, 4 gonads/19 nuclei; 6 h, 4 gonads/46 nuclei. The ZIM-3 and ZIM-2 analysis was performed on gonads fixed at 5 h post labeling (6 gonads/52 nuclei). Distances between foci were measured in 3D image stacks, as indicated. (B) Histograms depicting distance measurements between ZIM-3 foci and ZIM-3, ZIM-2 or HIM-8 foci. Distances were measured from the center of each major focus; distances below 0.8 μm were pooled into a single bin (asterisk). (C) Chart showing that the fraction of foci pairs that were not spatially resolved (“touching”) or that had inter-focus distance measurements below 1.2 μm were significantly higher for ZIM-3 – ZIM-3 foci pairs than for ZIM-3 – ZIM-2 pairs or ZIM-3 – HIM-8 pairs. P values are from two-tailed Fisher exact tests.

To simplify our analysis, we specifically considered only those nuclei in which all three assayed PCs were already paired, as indicated by a single HIM-8 (X-PC) or ZIM-2 (V-PC) focus and two separate (or immediately adjacent) ZIM-3 foci (corresponding to the PCs of chromosomes I and IV). We found that the fraction of foci pairs that were not spatially resolved (“touching”) or that had inter-focus distance measurements below 1.2 μm, were significantly higher for ZIM-3 – ZIM-3 foci pairs than for ZIM-3 – ZIM-2 pairs or ZIM-3 – HIM-8 pairs.

This observation of a bias toward shorter distances between ZIM-3-bound PCs suggests that the PCs of chromosomes that share the same PC-binding protein may spend more time in close proximity to each other than chromosomes that do not. This in turn raises the possibility that although PC-binding protein identity is not sufficient to enable pairing partner discrimination (Phillips and Dernburg 2006), transient grouping among chromosomes with matching PC binding proteins may nevertheless contribute to the chromosome sorting process in early meiotic prophase.

### Timing of SC assembly

We followed SC assembly using immunostaining of axis component HTP-3 and SC central region protein SYP-1 (Figures 6, 7). SYP-1 is first detected as a nuclear aggregate, typically colocalized with a non-axial pool of HTP-3 (Figure 6B). Such aggregates are a prominent feature of the leptotene stage (Figure 6C), as they were detected in nearly all assayed nuclei at the first time point analyzed for these features (3 h post injection) and rapidly disappeared as chromosome axes coalesced and SC assembly began upon transition from the leptotene to zygotene stage. In some nuclei in which HTP-3 axes were still in the process of coalescing, a single short stretch (or a few short stretches) of SYP-1 was detected in association with axial HTP-3. We observed a clear progression from nuclei with short SYP-1 stretches, to nuclei with an increasingly larger fraction of the autosomal HTP-3 axes associated with SYP-1, to nuclei where the full lengths of autosomal HTP-3 signals were SYP-1 associated, indicating completion of autosomal synapsis. Based on the stacked bar graph in Figure 6A and the event curves for autosomal SYP-1 pattern in Figure 6C, we can infer that the time window from initiation to completion of synapsis for all 5 pairs of autosomes is roughly 3.5 hours. This value is similar to, albeit somewhat shorter than, the 3.9-4.6 hour estimate for median time needed for completion of synapsis of all six chromosome pairs made by (Rog and Dernburg 2015) based on a combination of live imaging of SC assembly and computer modeling simulations.

**Figure 6.**
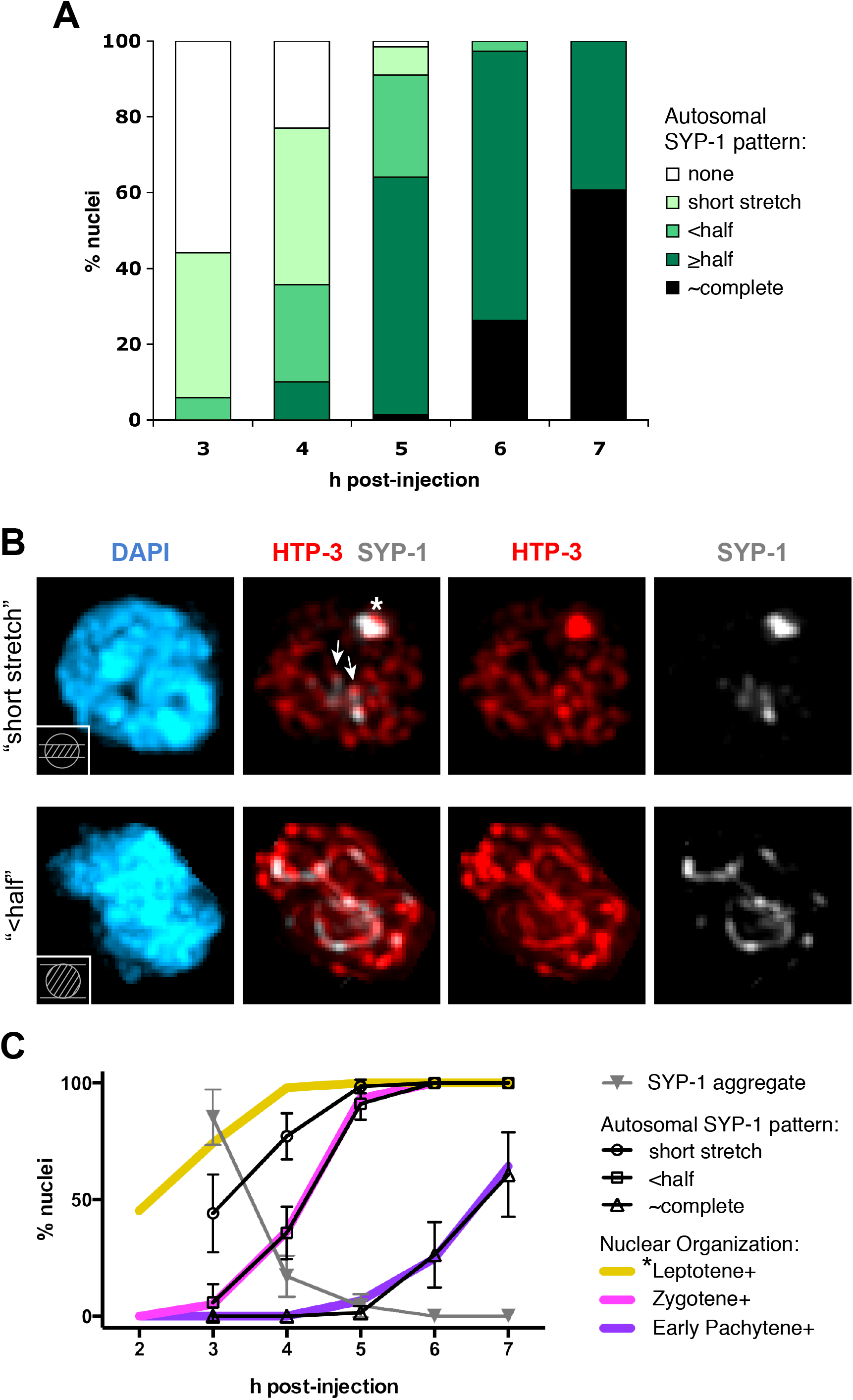
Timing of synapsis for autosomes. (A) Stacked bar graph indicating the percent of nuclei exhibiting the indicated patterns of SYP-1 localization associated with autosomes at each time point (in nuclei with S-phase labeled X chromosomes). For nuclei in the “short stretch” class (example shown in (B)), one or a few short stretches of SYP-1 were observed, together with a large SYP-1 aggregate not associated with chromosome axes; it is unclear whether such short stretches represent bona fide synapsis intermediates. The “<half” class comprises nuclei that lack a SYP-1 aggregate but have SYP-1 associated with less than half of the total length of the autosomal axes (marked by HTP-3 immunostaining). Numbers of gonads/ nuclei scored as reported in Figure 2 legend. (B) Sample images from the analysis quantified in (A); images are maximum intensity projections generated in Volocity from 3D image stacks (top, partial projection; bottom, full projection). Schematics in lower left corner of DAPI panels indicate the portion of the nucleus included in the projection. Top, nucleus from the “short stretch” category, showing two SYP-1 stretches that are clearly shorter than the length of a chromosome (arrows), plus a large SYP-1 aggregate (asterisk) that also contains (non-axis-associated) HTP-3 protein. Bottom, characteristic image from the “<half” class. (C) Cumulative distribution event curves for autosomal synapsis, indicating that it takes approximately 3 -3.5 hours from the onset of autosomal synapsis until synapsis is complete (see text). Also plotted is the actual frequency (not cumulative distribution) of SYP-1 aggregates present at the indicated time points, which indicates that SYP-1 aggregates are present in essentially all leptotene nuclei, but disappear at or soon after the onset of synapsis. All error bars indicate 95% CI.

**Figure 7.**
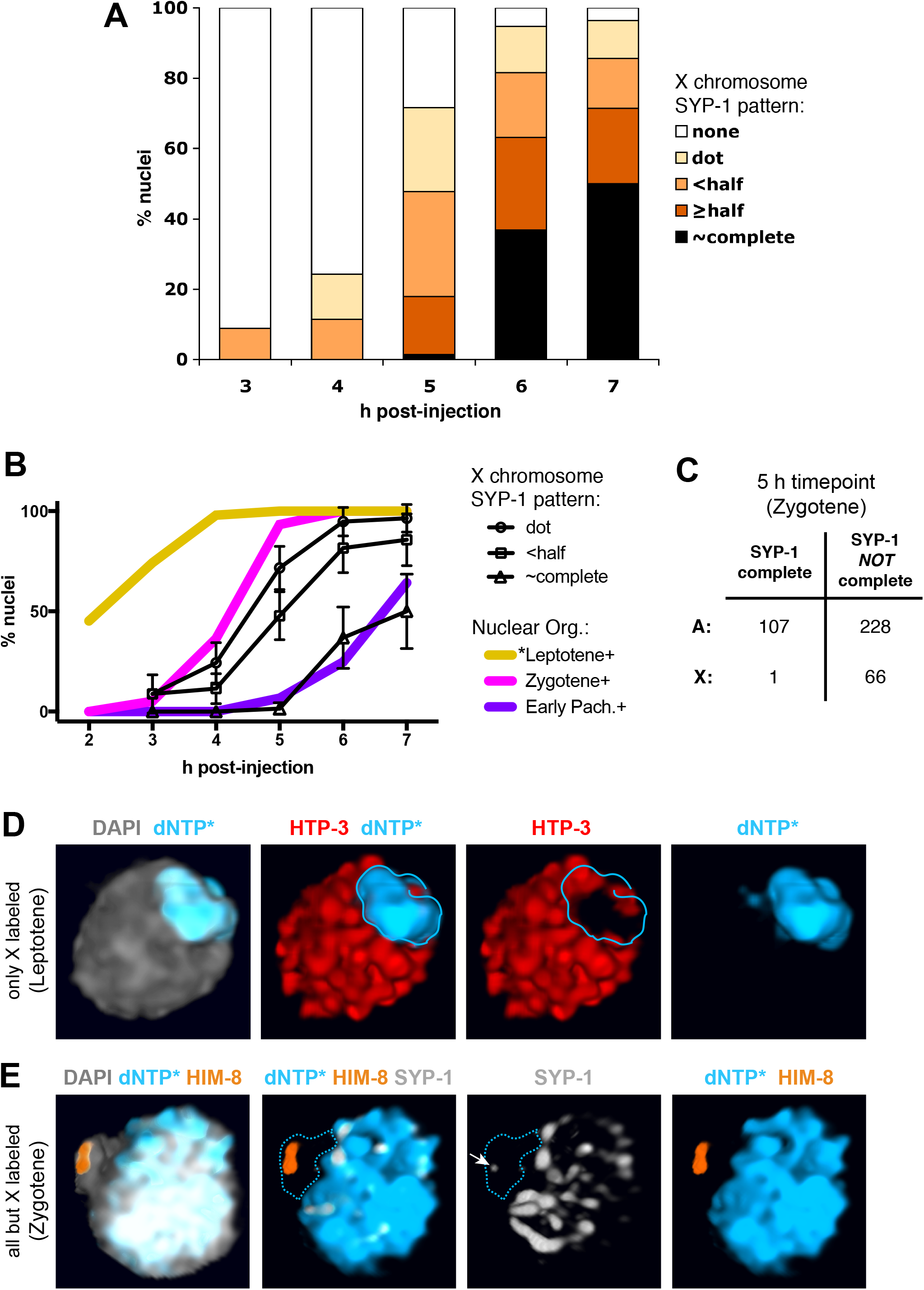
Timing of X chromosome synapsis. (A) Stacked bar graph indicating the percent of nuclei exhibiting the indicated patterns of SYP-1 localization associated with the X chromosome pair at each time point. In the “dot” class, the only SYP-1 associated with the X chromosomes is a small dot that has clearly not been extended at all (example shown in (E)); nuclei with any elongated SYP-1 stretch up to half the length of a chromosome pair comprise the “< half” class. Numbers of gonads/ nuclei scored as reported in Figure 2 legend. (B) Cumulative distribution curves for X chromosome synapsis (± 95% CI). (C) 2 x 2 contingency table depicting the incidence of fully synapsed autosomes relative to fully synapsed X chromosomes at the 5 h post injection time point, when essentially all of the nuclei were in the mid-zygotene stage. X chromosomes exhibiting complete synapsis were substantially underrepresented relative to autosomes ((p < 0.0001, two-tailed Fisher exact test). (D) Image panel depicting a leptotene nucleus 4 h post injection with S-phase labeled X chromosomes, immunostained for axis protein HTP-3. Whereas HTP-3 is associated with all autosomes in this nucleus, HTP-3 is not yet detected on the X chromosomes, suggesting that axis assembly on the X chromosomes is delayed relative to autosomal axis assembly. (E) Image panel of a zygotene nucleus from a “shadow time point” 8.5 h post injection, in which S-phase label had been incorporated into all autosomes during early S phase but not into the HIM-8-marked X chromosomes (as label had been depleted prior to the start of X replication). Whereas there is extensive SYP-1 associated with the autosomes in this nucleus, only a tiny SYP-1 dot is detected on the X (arrow). This image panel illustrates that later timing of SC assembly on the X chromosomes (relative to the autosomes) is not an experimental artifact resulting from incorporation of labeled nucleotides into the DNA. Images in (D and E) are from Volocity volume renderings of 3D data stacks encompassing whole nuclei.

As the X chromosomes are (by definition) specifically labeled in the synchronous cohorts of nuclei examined in our experiments, we were able to assess the timing of synapsis for that one specific chromosome pair (Figure 7). Based on the graphs in Figure 7A and B, we can generate upper and lower estimates for the average time from initiation to completion of synapsis of the X chromosome pair. If we assume that the appearance of a single SYP-1 “dot” on the X chromosomes is indicative of synapsis initiation, we can infer that it takes approximately 2.5 hours from initiation to completion of X-chromosome synapsis. Alternatively, using the appearance of linear stretches of SYP-1 as a more definitive indicator that synapsis has initiated results in an estimate of roughly 2 hours.

Notably, our 2- 2.5 hour estimate for the amount of time it takes to synapse the X chromosome pair is considerably longer than the 25-40 minute estimate of the time needed to complete synapsis of a chromosome pair reported by (Rog and Dernburg 2015). There are several possible explanations that, alone or in combination, may explain this discrepancy. First, features of our experimental design may have enabled detection of slow steps in the synapsis process not detected in the live imaging-based study. For example, the higher image quality and resolution afforded by fixed images, coupled with the use of an axis marker counterstain, may enable detection of earlier synapsis intermediates (“dots” or stretches <400 nm) that would not easily be detected or distinguished from background in live imaging; thus, the approach presented here may be capturing an inferred rate-limiting initiation step in SC assembly that was not captured in the live imaging. Similarly, it is possible that there may be a second slow step at the completion of SC assembly, and if so, this could only be detected by use of an axis counterstain. Differences in experimental temperature (20°C vs. ambient T ranging from 19 - 24°C for live imaging) may also be a relevant factor, as SC assembly is known to be a temperature-sensitive process (Bilgir *et al*. 2013). Finally, the rate of SC assembly for the X chromosome pair may be slower than the average synapsis rate for the five pairs of autosomes; in the prior live imaging study, the identities of the studied chromosomes were not known. An X vs. autosome difference in synapsis rate would not be surprising, as many features of chromosome organization in addition to the timing of replication are known to differ between the X chromosomes and the autosomes (Couteau and Zetka 2005; Jaramillo-Lambert *et al*. 2007; Jaramillo-Lambert and Engebrecht 2010; Kelly *et al*. 2002; Martinez-Perez and Villeneuve 2005).

### X chromosomes and autosomes differ in the timing of SC assembly

To address whether the X chromosomes and autosomes differ in the timing of SC assembly, we assessed the incidence of fully synapsed autosomes and fully synapsed X chromosomes at the 5 h time point, when essentially all nuclei were in the zygotene stage (Figure 7 C). This analysis revealed a striking difference between the X chromosomes and the autosomes: whereas 32% (107/335) of the autosome pairs in this cohort of nuclei were fully synapsed, only 1 of 67 X chromosome pairs exhibited complete synapsis (p < 0.0001, Fisher exact test). This finding clearly demonstrates that on average, the autosomes complete synapsis earlier than the X chromosomes. This may reflect a delay in initiation of synapsis and/or a slower rate of SC assembly for the X chromosomes relative to the autosomes.

During the early stages of meiotic prophase analyzed in our experiments (especially the 3- to 4-hour time points), the X chromosomes generally exhibit the weakest HTP-3 staining among all of the chromosomes in the nucleus (Figure 7D). This distinct appearance may reflect a delay in axis maturation caused by the late replication of the X chromosomes. We speculate that a difference in structure and/or subunit composition between X chromosome axes and autosomal axes may in turn affect the timing of onset and/or rate of SC assembly. We note that this probably does not reflect the X chromosomes having an inherently lower affinity for SYP proteins, however, as SCs are detected preferentially on the X chromosomes when the SYP-1 protein is present in limiting amounts (Hayashi *et al*. 2010).

### Concluding remarks

The detailed and comprehensive analysis of the timing of and relationships among key early events in the *C. elegans* meiotic program, summarized in Figure 8, represents an important contextual framework that can facilitate and augment future research. Further, it provides strong validation for the use of surrogate staging markers (such as specific DAPI morphologies) and other temporal landmarks that can serve as linchpins for cytological analyses of meiotic mechanisms; this will be especially useful for imaging experiments where not all relevant features can be visualized simultaneously. Conversely, it emphasizes the efficacy of S phase labeling of X chromosomes as a synchrony marker that enables temporal staging in mutants where normal staging markers are absent. Finally, our findings demonstrate the capacity of this approach to reveal aspects of the meiotic program (*e.g*. X vs. autosome differences in synapsis timing, or preferential proximity of PCs sharing the same PC-binding protein) that are not readily detectable by analyses using spatially-defined zones for feature quantitation. We hope that this work will serve as a valuable resource for researchers investigating the mechanisms of meiosis in the *C. elegans* system.

**Figure 8.**
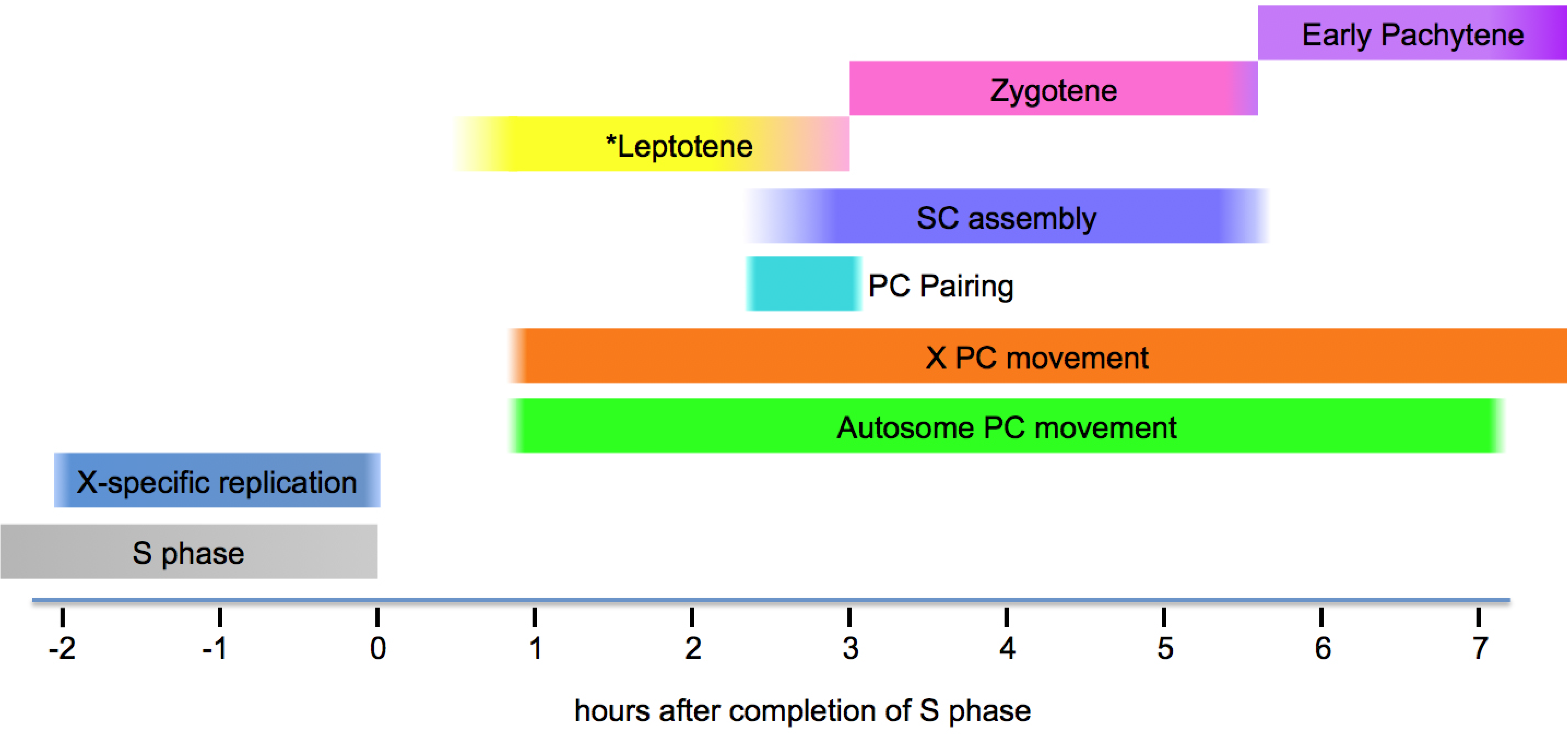
Summary schematic depicting onset and duration of key meiotic prophase events in *C. elegans*. For the meiotic prophase events and stages depicted, the time of onset for each event/stage reflects the time at which 50% of nuclei in the scored cohort had initiated that event or reached that stage. Likewise, time of completion reflects the time at which 50% had completed that event or transitioned to the next stage. Times are relative to the end of S phase (indicated as t=0), which corresponds to the 1 h post injection time point.

Movie S1. Movie of Cy3-labelled chromosomes(red) and ZYG-12::GFP(green) in a field of germ cell nuclei from a partially extruded gonad imaged at 3h 20m post injection (orientation: R to L = distal to proximal, flipped relative to the orientation shown in Figure 3A). Movie was rendered using projections of image stacks (3 optical sections; 1.5 μm step size) acquired every 10 s over a 2 min time period. Video plays at 7 fps (70x speed).

Movie S2. Movie of Cy3-labelled chromosomes (red) and ZYG-12::GFP(green) in a field of germ cell nuclei from a partially extruded gonad imaged at 8 h post injection (orientation: lower L to upper R = distal to proximal, rotated approximately 45 degrees counterclockwise relative to the orientation shown in Figure 3A). Movie was rendered using projections of image stacks (3 optical sections; 1.5 μm step size) acquired every 10 s over a 2 min time period. Video plays at 7 fps (70x speed).

## Acknowledgements

We thank the Caenorhabditis Genetics Center (funded by NIH Office of Research Infrastructure Programs P40 OD010440) for strains, A. Dernburg for antibodies, and C. Akerib, B. Roelens and A. Woglar for comments on the manuscript. This work was supported by an American Cancer Society Postdoctoral Fellowship and a Beckman Senior Research Fellowship to SME and by an American Cancer Society Research Professor Award (RP-15-209-01-DDC) and NIH grant R01GM53804 to AMV.

